# Hidden in plain sight: the invasive macroalga *Caulerpa prolifera* evades detection by environmental DNA methods due to negligible shedding

**DOI:** 10.1101/2023.07.31.551316

**Authors:** Tanner Waters, Kylie Langlois, Zack Gold, Susanna Theroux, Robert A. Eagle

## Abstract

The macroalgae *Caulerpa prolifera* is considered an invasive species in many environments and can colonize large patches of seafloor, reduce native species, and alter ecosystem functioning. Environmental managers need a rapid and cost-effective monitoring tool for tracking the spread of this invasive species. We developed a digital PCR assay for detection of *C. prolifera* from environmental DNA seawater samples. We demonstrate, in both field and laboratory experiments, that the invasive algae *C. prolifera* is undetectable in practical applications of eDNA due to its minimal shedding. To test why, we conducted tank-based shedding experiments for two California invasive algae species, *C. prolifera* and *Sargassum horneri*. Copy numbers of *C. prolifera* eDNA detected in the experimental tanks were found to be two orders of magnitude lower than *S. horneri*. A meta-analysis of steady state eDNA produced by aquatic organisms reported in the literature show *C. prolifera* to have the lowest recorded steady state concentrations of eDNA in the water column. We attribute *C. prolifera* low eDNA shedding to its unique biology as a unicellular, multinucleate, macroscopic siphonous algae which reduces the possible modes of eDNA release compared to multicellular organisms. Our results highlight the value of benchmarking and validating eDNA surveys in both field and laboratory settings and potential limits of eDNA approaches for some applications. These results also emphasize the importance of organismal physiology in eDNA shedding rates, variations in mechanisms of eDNA shedding between organisms, and characterizing shedding rates for accurate interpretation of eDNA results.

## Introduction

Invasive species are a threat to global marine biodiversity (Molnar et al., 2008). When non-native species are introduced to a new environment, they can rapidly colonize the area because of their quick reproduction time, lack of natural predators, ability to outcompete native species, or a combination of all three. This causes both direct and indirect impacts to local ecosystems. Due to their ability to outcompete native species, invasives often alter local biodiversity, impact ecosystem structure and reduce functional ecosystem services (Pimentel et al., 2000). Globally, marine invasive species have caused the economy an estimated $345 billion in damages (Cuthbert et al., 2021). This threat has only continued to rise in recent decades (Seebens et al., 2020) with the increase in globalized shipping, aquaculture, and accidental release (Bax et al., 2003; De Silva et al., 2009; Hulme, 2009). For these reasons, early detection and eradication before spread is a top priority for environmental managers (Larson et al., 2020).

One invasive species of particular concern is *Caulerpa taxifolia. C. taxifolia* is one of the top 100 worst invasive species (Global Invasive Species Database, 2023) and is designated on the US Federal Noxious Weed List due to its history of overtaking marine ecosystems. It received this level of scrutiny because in the first 16 years since its introduction off the coast of Monaco in 1984, it grew to cover nearly 131 km^2^ of Mediterranean coastline (Meinesz et al., 2001). The alga was first seen in 2000 in California in Carlsbad and Huntington Harbor, California (Jousson et al., 2000) with DNA barcoding of the tissue showing that it likely originated from an aquarium store (Jousson et al., 2000). It took nearly six years and seven million dollars (USD) to eradicate *C. taxifolia* from California (Merkel & Associates, 2006). In 2021, the first known case of *Caulerpa prolifera* was discovered off the West Coast of the United States in Newport Bay, CA (NOAA Fisheries). *C. prolifera* has been observed to stunt ecosystem services, reduce native biodiversity and significantly decrease species richness compared to native seagrass meadows (Parreira et al., 2021). These impacts and the species’ relative ease in spreading made it a top priority for eradication efforts by local marine managers.

Conventional survey techniques to identify *C. prolifera* involve divers visually scanning the seafloor. This presents an issue in Newport Bay where the turbidity makes for poor visibility and identifying *Caulerpa* amongst dense eelgrass allows divers to miss fragments. Survey efficacy through the use of artificial *C. prolifera* released in the bay found that nearly 20% of the fake fragments were never recovered (Owens, 2021). To aid in detecting *C. prolifera*, an environmental DNA (eDNA) approach was proposed. Environmental DNA is an emerging surveying tool that captures shed/excreted DNA from an organism by filtering an environmental water sample (Thomsen and Willerslev, 2015). Due to eDNA’s methodology of capturing free floating DNA from the water column, it has been shown to better detect rare and cryptic species and outcompete conventional survey techniques in the field (Thomsen and Willerslev, 2015; Fediajevaite et al., 2021). This potentially makes it a valuable approach for surveys aimed at detecting invasive species where early identification and removal are paramount to their eradication (Larson et al., 2020).

California has dealt with another invasive macroalgae for nearly 20 years. *Sargassum horneri* (Devil’s weed) is an alga native to Eastern Asia and was introduced to the West Coast of North America in 2003. In the 20 years since its introduction, its range has expanded from Baja California, Mexico to Point Conception, California including the Channel Islands (Marks et al., 2017). *S. horneri* often forms large mats off the coast that are anchored to rocks. Researchers working on *S. horneri* have shown that removal techniques are most effective in culling population size and density if the size of removal is enough to reduce propagule supply (Marks et al., 2017). Environmental DNA could be used for early detection of *S. horneri* prior to introduction via ballast water, for detection in areas that are difficult to survey, or for detection of small populations before they’re too large for eradication efforts. Benchmarking an *S. horneri* assay and its shedding rates is needed to validate eDNA as an appropriate monitoring method for these applications.

Our study looks to use environmental DNA to work alongside conventional survey techniques for the tracking and eradication of *Caulerpa*. We report the creation of a novel dPCR assay for the *in-situ* identification of *Caulerpa prolifera* and characterize the first algal eDNA shedding rates in the literature for *Caulerpa prolifera* and *Sargassum horneri* to benchmark this methodology as a monitoring tool for invasive algal species.

## Materials and Methods

### Assay design

To create our eDNA assay, we downloaded reference sequences of *Caulerpa prolifera* from GenBank (Bensen et al., 2015) (https://www.ncbi.nlm.nih.gov/genbank/). Sequences were aligned in Geneious 2019.2.3 (https://www.geneious.com) and potential primer/probe sets were created using Geneious’ design new primer/probes feature with guidelines based on Klymus et al 2020. Our primers were created for the ‘internal transcribed spacer’ or ITS gene based on previous work which has used the ITS for *Caulerpa* sp. phylogenetics (Kazi et al., 2013; Stam et al., 2006). Primer specificity was tested *in-silico* using EcoPCR (Ficetola et al., 2010) and showed species-specific *Caulerpa prolifera* amplification.

### Field testing

A *Caulerpa prolifera* patch was found in China Cove, Newport Bay via scuba diving surveys and was roughly one foot in diameter and contained ∼20 fronds. We sampled seawater on June 30th, 2021, directly above the bed (33.596406, -117.879731), and then above the sea floor 5m, 10m, 50m 100m, and 500m downstream from the Caulerpa patch, employing the eDNA collection method of Curd et al. 2019. First, we collected seawater samples directly above the patch using divers as to not disturb the *Caulerpa*. Samples taken away from the patch were then collected using a 5L niskin bottle. From the niskin, we transferred one liter of seawater to a Kangaroo enteral feeding bag in triplicate. We immediately filtered 1L of seawater through a sterile 0.22 μm Sterivex cartridge filter (MilliporeSigma, Burlington, MA, USA) through gravity filtration. We capped the filters and stored them on dry ice during sampling until we returned to the lab where they were stored at -20°C. Additionally, we filtered one liter of Milli-Q water through the same process for a negative field control (Goldberg et al., 2016). Tissue samples from the patch were taken for species verification.

### Experimental design of shedding experiment

We tested the shedding rates of two California invasive macroalgae, *Caulerpa prolifera* and *Sargassum horneri*. We ordered the *Caulerpa prolifera* from an online aquarium store and divers from Cabrillo Aquarium identified and collected the *Sargassum horneri* off the coast of San Pedro, CA. The algae were left to acclimate in tanks for two days before the start of the experiment. We filled three replicate tanks per species with 20L of DI water and 36 g/L of Instant Ocean sea salt for aquariums. Wet weights of the algae were measured and recorded before they were added to the tanks. We kept the tank water at ambient room temperature in the lab (20 ± 1 °C) and exposed to natural, indirect sunlight through the window. An additional tank containing only artificial seawater was used as a control.

We took samples before the addition of the species (hour 0) and then subsequently at 1, 2, 4, 8, 12, 24, 48, 72 and 96 hours after they were added. We added 23.99g, 24.44g, and 23.39g of *Caulerpa* and 20.47g, 23.49g, and 22.36g *Sargassum* into their respective first, second and third tanks. When collecting the samples, we stirred the tank gently with a sterile stirrer for a well-mixed sample and then collected 1L of tank water into a Kangaroo enteral feeding bag. This 1L bagged sample was then filtered onto two sterile 0.22 μm Sterivex cartridge filters (MilliporeSigma, Burlington, MA, USA) running 500mL through each via gravity filtration to avoid filter clogging. We stored the filters at -20°C until they were extracted the following day. After each sample collection time point, we immediately refilled the tank with 1 L of sea water from a carboy so as to maintain consistent volume within the small tanks. At each time point, we collected water from the control tank and carboy in the same manner to test for contamination.

For the *Caulerpa* tanks, we took additional environmental RNA samples at hour 0 and hour 96 in the same manner described above for the environmental DNA samples. We stored the filters at - 80°C until they were extracted within a couple weeks.

### DNA/RNA Extraction

All eDNA samples we extracted from the Sterivex cartridge using a modified DNeasy Blood & Tissue Kit protocol (Qiagen Inc., Germantown, MD) optimized for increased eDNA yield (Spens et al., 2017). Sterivex filters were incubated at 56°C overnight with 720μl of ATL buffer and 80μl of proteinase K. After incubation, equal parts AL buffer and ice-cold molecular grade ethanol were added to the ATL buffer/ proteinase K mixture and spun through a spin column. AW1 and AW2 buffers were added to wash the columns. The DNA was eluted using 100μl of AE buffer and stored at -20°C.

Environmental RNA was extracted using a RNeasy Power Water Kit (Qiagen Inc., Germantown, MD) following the manufacturer’s protocol modified for sterivex filters as shown above.

### Assay Conditions

Our droplet digital PCR assay amplifies a 106 bp amplicon and uses the forward primer 5’-TGGCGCTATGTAATGTTGATGTTG-3’, reverse primer 5’-GCAATTCGCAACACCTTTCGTA-3’ and probe 5’-56-FAM-CGGTTCCCGTGTCGATGAAGGACG-3IABkFQ-3’. We ran an annealing temperature gradient to find the temperature with the best amplification and greatest difference between positive and negative droplet fluorescence amplitudes. Based on this, cycling conditions for the dPCR were 95°C for 10 minutes, 40 cycles of 95°C for 30 seconds and 58°C for 60 seconds, 98°C for 10 minutes and a 4°C hold indefinitely. Mastermix concentrations were: 14.4 μl of 4x ddPCR Multiplex Supermix (Bio-Rad, Hercules, CA), 0.5184 μl of 100uM forward primer, 0.5184 μl of 100uM reverse primer, 0.144μl of 100uM probe, 20.4192μl of water and 12 μl of sample. This mix was partitioned into duplicate replicates of 22μl and added to a 96 well plate. The reaction mixture was combined with Bio-Rad Droplet Generation Oil (20 μl reaction mixture + 70 μl oil) and partitioned into nanodroplets via microfluidics in the Automated Droplet Generator (Bio-Rad). This resulted in a total nanodroplet volume of 40 μL, which was transferred to a standard 96-well PCR plate for amplification using a multichannel pipettor. The plate was heat sealed with pierceable foil using a PX1 PCR plate sealer (Bio-Rad) and PCR amplification was carried out in a S1000 thermal cycler (Bio-Rad, ramping speed at 2°C per second). After PCR, the plate was read by the Bio-Rad QX200 Droplet Reader and analyzed using the Bio-Rad QX Manager (v.1.2 or v.2.0) software. The *Caulerpa* eRNA samples followed the same steps except with a mastermix of 12μl of MM, 4.8μl of RT, 2.4μl of 300mM DTT, 0.5184 μl of 100uM forward primer, 0.5184 μl of 100uM reverse primer, 0.144μl of 100uM probe, and 15.6192 μl of water and one hour at 50°C prior to the cycling conditions for reverse transcription to occur.

*Sargassum horneri* primers were chosen from Hamaguchi et al., 2022. We ran an annealing temperature gradient on the primers which showed a 56°C annealing temperature to be the optimum temperature. All other assay conditions were the same as described above for the *Caulerpa* assay.

### Data Analysis

Following recommendations in Cao et al. 2015 and Steele et al. 2018, a minimum of two reactions and a total of ≥10,000 droplets per reaction were generated per sample; samples that failed to meet the droplet requirement were reanalyzed. At least six no template control (NTC, RNA/DNA-free water; UltraPureTM, Life Technologies, Carlsbad, CA, USA) reactions were run per assay. NTC samples were required to contain less than 3 positive droplets. Two positive control reactions were included per assay. When inhibition was observed, or samples exceeded the upper limit of quantification, these were diluted with RNA/DNA-free water and reanalyzed.

Based on the concentrations from the ddPCR software, we back calculated the tank concentration of DNA using the following code https://github.com/kylielanglois/SCCWRP/blob/main/ddPCR/ddPCR_autofill_clean.R. The ddPCR output in copies/μL of reaction were converted to copies/μL in the filter and then converted to copies/μL of tank water. Since replicates came from the same 1L bag, they were averaged together to account for larger particles that were unevenly distributed between the two filters.

We then calculated the steady state concentration per gram of body mass using the equation from Sassoubre et al., 2016. Briefly, *V*dC/dt = S-kCV* where V is the tank volume in liters, C is eDNA concentration, t is hours, S is the shedding rate and k is the first-order decay rate constant per hour. At steady state, dc/dt= 0 so the shedding rate/decay rate constant would equal the concentration of eDNA multiplied by the tank volume. Since our experiment did not measure the decay rate constant, we can solve for the steady state concentration per gram of body mass by using the 96-hour concentration when our tanks reached steady state. We then compare steady state concentration per gram of body bass across other previously reported values for other species (Andruszkiewicz Allan et al., 2020; Kwong et al., 2021; Maruyama et al., 2014; Nevers et al., 2018; Plough et al., 2021; Sansom and Sassoubre, 2017; Sassoubre et al., 2016; Wilder et al., 2023). When shedding and decay rates for multiple conditions in a given study for a single species was reported, we report both the lowest and highest reported steady state to show the range. All values shown are from other tank-based, single species shedding experiments allowing for comparable results within the meta-analysis.

## Results

### Field Sampling

We successfully amplified extracted *Caulerpa prolifera* tissue DNA collected from the field invasion using our Caulerpa-specific primer set. A serial dilution of tissue DNA showed congruent copy number and minimal dPCR rain which indicated sufficient PCR efficiency. Despite this, none of the field eDNA samples found any detectable amounts of *Caulerpa prolifera* DNA, including the samples taken directly above the visually observed patch. Additional replicate field samples taken directly above the *Caulerpa* patch were tested for inhibition using a serial dilution and a Qiagen DNeasy PowerClean Pro Cleanup Kit and similarly showed no amplification of *Caulerpa prolifera* DNA.

### Tank-based Experiment

The *Caulerpa prolifera* and *Sargassum horneri* in the tank experiments both produced quantifiable eDNA (Supplemental Table 2; Supplemental Table 3). *Caulerpa* was characterized by a sharp increase in initial eDNA concentration in the tank followed by a steady plateau. *Sargassum* instead saw a general increase in eDNA tank concentration followed by a similar plateau. Our sampling method of filtering 1L and replacing it with 1L of water may have diluted the concentrations by 5%, which would have no bearing on the final interpretation of the results. Both tanks reached steady state by 96 hours where no observable change in the concentration of tank eDNA was seen. For the 20g samples of algae in each tank, this steady state equates to roughly 10^4.5^-10^5^ copies of DNA/ L of tank water for *Caulerpa* and 10^7^ copies/L for *Sargassum* (Fig. 1). This equates to a nearly 100-315x greater amount of *Sargassum* eDNA concentration per gram of biomass compared to *Caulerpa*. All tank controls and PCR controls were blank.

**Fig. 1.**
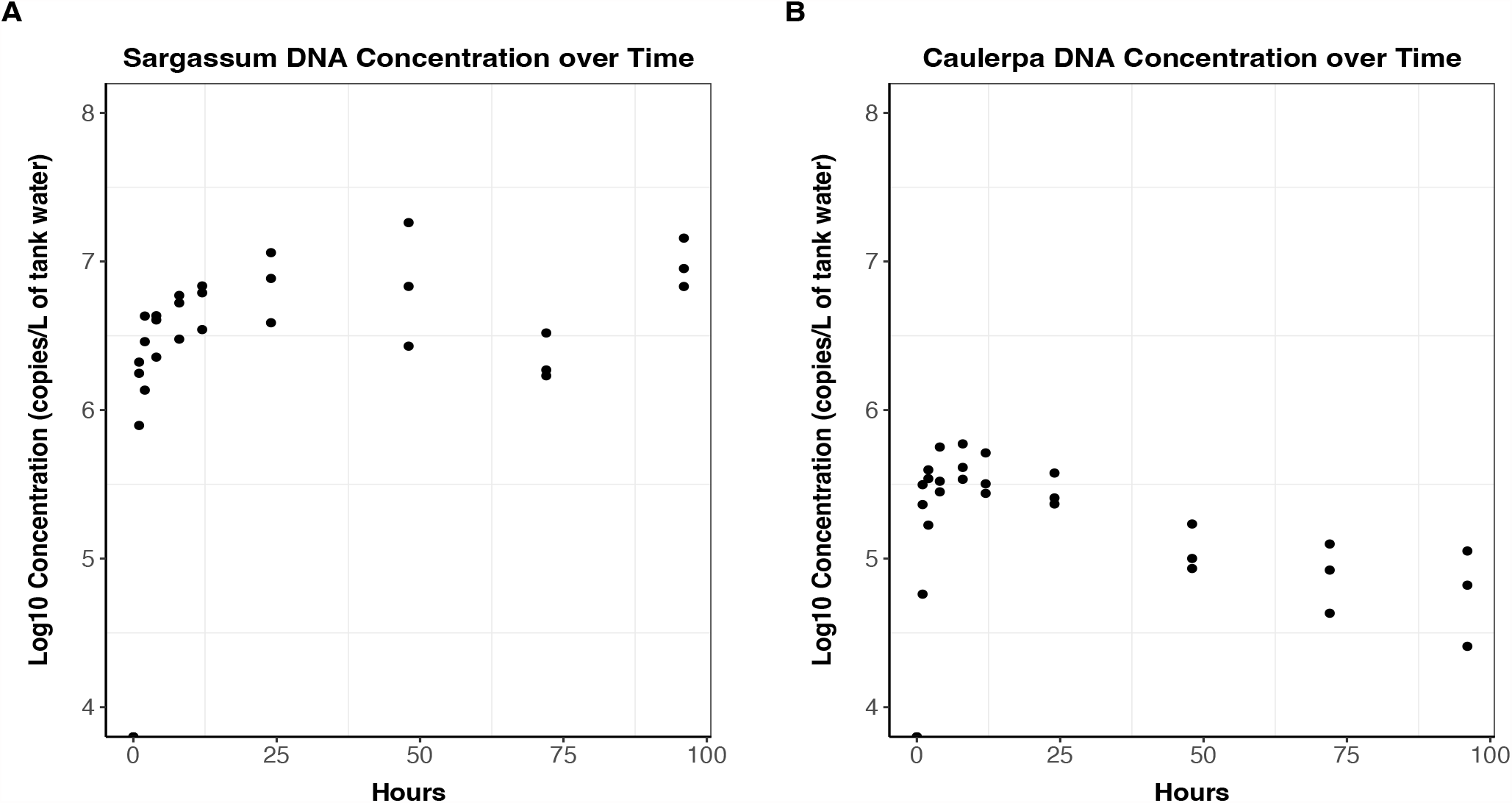
Plots of the tank eDNA concentrations over time in log form. Sargassum shows an initial jump in eDNA copies/L and then a steady plateau after 24 hours. Caulerpa exhibits an initial spike in concentration before decreasing and leveling off after 48 hours. Steady state for both was reached at the 96 hour time point.

### Environmental RNA

The *Caulerpa* environmental RNA (eRNA) samples showed detectable, but minimal, RNA copy numbers per tank. At 96 hours, eRNA was found to be 345-1630 copies/L of tank water with one replicate being under the ddPCR limit of quantification. For comparison, eDNA concentrations at 96 hours were roughly 70-75x as much as eRNA.

### Steady State

Steady state copies of DNA per gram of biomass was calculated for the available species in the literature, spanning over 7 orders of magnitude (Supplemental Table 1). *Caulerpa prolifera* had the lowest steady state concentration while *Sargassum horneri* was one of the median reported values. (Table 1; Fig. 2)

**Table 1.** Steady state concentrations of environmental DNA from existing literature. When shedding and decay rates for multiple conditions in a given study for a single species was reported, the lowest and highest reported steady state was calculated to show the range.

| Citation | Species | Steady State<br>(Copies/g) |
| --- | --- | --- |
| Andruszkiewicz Allan et al., 2020 | *Chrysaora spp.* (Sea Nettle) | 2.04 x 10<sup>11</sup> |
| Sassoubre et al., 2016 | *Sardinops sagax* (Pacific sardine) | 1.62 x 10<sup>11</sup> |
| Plough et al., 2021 | *Acipenser oxyrhynchus oxyrhynchus* (Atlantic sturgeon) | 1.43-1.58 x 10<sup>11</sup> |
| Andruszkiewicz Allan et al., 2020 | *Aurelia aurita* (Moon jelly) | 9.16 x 10<sup>10</sup> |
| Andruszkiewicz Allan et al., 2020 | *Fundulus heteroclitus* (Mummichog) | 8.73 x 10<sup>10</sup> |
| Sassoubre et al., 2016 | *Scomber japonicus* (Pacific Chub mackerel) | 6.19 x 10<sup>10</sup> |
| Andruszkiewicz Allan et al., 2020 | *Palaemon spp.* (Grass shrimp) | 2.14 x 10<sup>10</sup> |
| Sassoubre et al., 2016 | *Engraulis mordax* (Northern anchovy) | 1.14 x 10<sup>10</sup> |
| **This study** | *Sargassum horneri* (Devil's Weed) | 5.77 x 10<sup>6</sup> - 1.23 x 10<sup>7</sup> |
| Maruyama et al., 2014 | Juvenile *Lepomis macrochirus* (Bluegill sunfish) | 1.12 x 10<sup>7</sup> |
| Nevers et al., 2018 | *Neogobius melanostomus* (Round goby) | 5.17-5.81 x 10<sup>6</sup> |
| Wilder et al., 2023 | Juvenile *Esox masquinongy* (Muskellunge) | 2.7-3.45 x 10<sup>6</sup> |
| Maruyama et al., 2014 | Adult *Lepomis macrochirus* (Bluegill sunfish) | 2.6 x 10<sup>6</sup> |
| Kirtane et al., 2021 | *Paralichthys dentatus* (Summer flounder) | 5.06 x 10<sup>5</sup> – 3.28 x 10<sup>6</sup> |
| Kirtane et al., 2021 | *Pseudopleuronectes americanus* (Winter flounder) | 5.24 x 10<sup>5</sup> – 1.32 x 10<sup>6</sup> |
| Kirtane et al., 2021 | *Centropristis striata* (Black sea bass) | 3.01 x 10<sup>5</sup> – 1.64 x 10<sup>6</sup> |
| Wilder et al., 2023 | Larval *Esox masquinongy* (Muskellunge) | 2.44 x 10<sup>5</sup> - 1.21 x 10<sup>6</sup> |
| Kwong et al., 2021 | *Acanthaster cf. solaris* (Pacific crown-of-thorns sea star) | 3.76 x 10<sup>5</sup> |
| Sansom and Sassoubre, 2017 | *Lampsilis siliquoidea* (Freshwater mussel) | 5.26 x 10<sup>4</sup> - 4.91 x 10<sup>5</sup> |
| **This study** | *Caulerpa prolifera* | 2.1-9.38 x 10<sup>4</sup> |

**Fig. 2.**
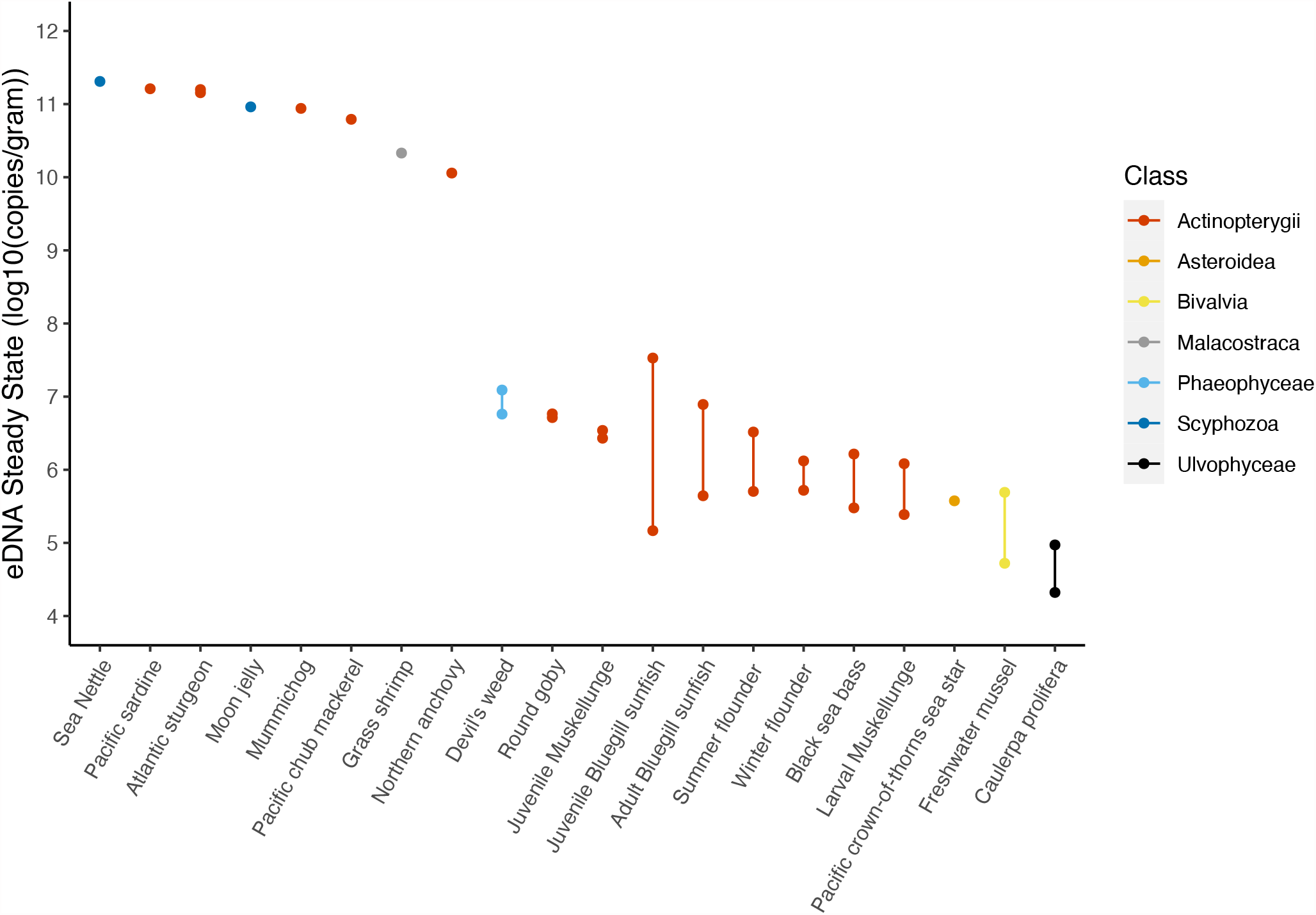
Log10 conversion of the steady state concentration of eDNA by species and class. Where variable steady states were reported between experiments, we plot the lowest and highest rates reported and the range in values is indicated by the bar linking two points. Those with one steady state rate only reported one shedding and decay rate.

## Discussion

Our results demonstrate the importance of lab and field validating eDNA methods prior to their adoption as a monitoring technique. We created a novel eDNA assay that is able to amplify *C. prolifera* DNA. Despite this, we were unsuccessful in identifying *C. prolifera in-situ* over a known patch of the algae. Tank-based experiments demonstrate that *C. prolifera* has the lowest observed steady state eDNA concentration of any reported species. In contrast, *Sargassum horneri* assay shows promise as an invasive monitoring tool due to its higher steady state concentration. Our results have implications on the use of eDNA in the field of invasive species monitoring and on our understanding of eDNA shedding mechanisms.

### *Caulerpa* sheds negligible amounts of DNA

This study shows *C. prolifera* to have the lowest recorded eDNA steady state concentration of any currently reported species. The low concentration of *Caulerpa* eDNA can best be attributed to the algae’s unique organismal and cellular biology. The genus *Caulerpa* is home to some of the largest single celled organisms in the world (Jacobs, 1994). *C. prolifera* is a multinucleated single celled macroalgae which spreads primarily through asexual reproduction (Jacobs, 1994). This means that it lacks conventional modes of eDNA release including shed cells via feces, urine, blood, gametes, hair, etc., which account for a considerable amount of total eDNA release for other species (Klymus et al., 2015; Thomsen and Willerslev, 2015). As a single-celled organism, it itself is not shedding cells into the water but is at the same time, it is much too large of a cell to be captured within our water samples as is the case for bacteria and phytoplankton species. This leaves only a few potential modes of eDNA shedding, namely cellular leakage of mitochondria or free-floating DNA. Previous work has demonstrated that cellular leakage accounts for only a small portion of total environmental DNA (Zhoa et al., 2021). Further limiting DNA shedding is *Caulerpa’s* thick cell wall surrounding the algae which prevents regular shedding of cellular material into the water column (Jacobs, 1994), providing substantial support for the hypothesis that the unique physiology of *Caulerpa* contributes to low shedding rates.

In addition to *Caulerpa*’s unique single celled physiology, previous studies have shown dramatic differences in gene expression across the organism despite being a single cell, likely explaining the unique morphology of fronds, rhizomes, etc., and suggesting additional cellular mechanisms to limit DNA and RNA activity and transport (Arimoto et al., 2019; Ranjan et al., 2015). It is currently unknown what mechanisms allow for such differentiation within the organism despite shared cytoplasm. However, one potential explanation may be a high degree of RNAse and DNAse activity which would limit the spread of transcription and translation to specific regions of the organism, allowing for the substantial phenotypic differentiation observed across the cell. Such a mechanism may also act to reduce the amount of free-floating DNA available within the organism and thus impact shedding rates via cellular leakage. Together, these factors strongly suggest that the unique physiology and morphology of *Caulerpa* contribute to low shedding rates. Unfortunately, our results indicate that this invasive species is uniquely equipped to evade detection through eDNA surveying, indicating the need for alternative detection methods.

We note that the other reported values from previously reported shedding and decay experiments also underestimate the amount of eDNA release that occurs *in-situ*. During tank-based trials, animal species are restricted from food before and during the experiment to minimize the amount of eDNA introduction to the tanks from sources such as feces. This means in the wild, when species have access to food and have higher metabolic rates, expected eDNA release rates are higher. Previous work has demonstrated that sea stars in tank experiments that were given food released roughly 7x more eDNA than when not given food, strongly supporting this hypothesis (Kwong et al., 2021). Thus, given that the majority of steady state concentrations were generated from heterotrophic species, we expect their relative eDNA shedding rates to be even higher than the photosynthetic *Caulerpa*, contributing to the low detection rates of this invasive species.

Furthermore, a large number of aquatic plant and animal species also introduce eDNA through the release of gametes during spawning events. However, *Caulerpa’s* predominantly asexual mode of reproduction limits shedding rates compared to broadcast spawning organisms (Smith and Walters, 1999).

We note that *Caulerpa* steady state concentrations at 96 hours were 5-20% of the maximum concentration (Fig. 1). However, previous tank experiments results show an initial spike in DNA concentration as a result of stress to the organism (Klymus et al., 2015; Nevers et al., 2018) followed by a decline in production. All values used in our comparisons of steady state concentrations were reached within 24-48 hours into their respective experiments so as to avoid any differential physiological effects of initial stress influencing our comparisons. Thus, we are confident that values presented for *Caulerpa* in this study more appropriately captures the steady state concentration it would experience in the wild.

Additionally, the steady state values calculated were normalized by grams of biomass so that the values were comparable across taxa. This metric undervalues the difficulty in detecting *Caulerpa in-situ* compared to other species in this list. The next lowest mean steady state concentrations are from the freshwater mussel and Pacific crown-of-thorns sea star (Sansom and Sassoubre, 2017; Kwong et al., 2021). These species weigh roughly 100g and 3000g, respectively, whereas a single *C. prolifera* frond weighs just a fraction of a gram. A colony of *Caulerpa* that consists of 100-3000g of biomass would make environmental DNA as a tool unnecessary because the patch, likely to be multiple square meters in size depending on its density, would be visible to conventional visual surveys.

### eDNA as a tool to track *Sargassum*

Previous *S. horneri* research found that removal efforts have considerable challenges to success once the alga has been established (Marks et al., 2017). This emphasizes the importance of early detection to the protection of our coastal ecosystems. This study calculated *Sargassum horneri*’s steady state concentration to be roughly 10^6^ which placed it within the middle range compared to fish and invertebrates. We demonstrate here that because of its relatively high steady state concentration, and large biomass in the wild, *Sargassum* is an ideal invasive candidate for environmental DNA detection as demonstrated in previous studies via metabarcoding (Ely et al. 2021 and Gold et al 2022). Specifically, the use of this ddPCR assay in areas with low abundance before species establishment, environments of high turbidity and low visibility, locations that are difficult to dive in, and in ballast water of ships would allow for higher sensitivity monitoring and earlier intervention. State and federal agencies such as CDFW, NOAA, USGS, and USFWS that are tasked with monitoring and stopping invasive species would particularly benefit from the use of eDNA to monitor *Sargassum horneri* populations.

### Implications for environmental DNA studies

The results of this study show a multiple order of magnitude difference in species’ eDNA steady state concentrations. The differential steady state values highlight the influence various eDNA release modes play in detection probabilities and the difficulty in ascribing quantitative metrics to eDNA data between species. Notably, fish species show a significant range of steady state concentrations from 10^6^-10^11^ copies/gram. Fish with a higher steady state concentration have higher probabilities of being detected in the wild and would therefore be overrepresented in environmental DNA surveys, all else being equal, compared to fish that have 5 orders of magnitude less concentration. As the field of eDNA moves to be more quantitative, relative abundances such as the eDNA index which use a double proportion to convert raw read counts, will be important in accounting for uneven shedding rates (Kelly et al., 2019). These results suggest that relative shedding rates operate on similar orders of magnitude as amplification efficiencies and thus controlling for both biases will likely be critical for deriving quantitative metabarcoding approaches (Shelton et al. 2022). Our meta-analysis here also highlights the limited number of non-fish species in eDNA shedding and decay experiments. As eDNA aims to become a holistic monitoring tool for biodiversity, characterizing shedding and decay relationships across a broad diversity of taxa and not just those that are commercially important, will be crucial to understanding the applicability of this methodology for biodiversity monitoring efforts.

Previous work has argued that eDNA in its current state works best as a complement to conventional survey techniques (Bohmann et al 2014; Kelly et al. 2017). In the case of invasive species, eDNA can aid in early detection of areas of concern while there is always value for “boots on the ground” confirmation, especially when there are significant management implications (Gold et al. 2022). A strong advantage of eDNA is the ability to reduce the complexity of the field logistics by narrowing the range of visual surveys and the time it would take to complete them. However, our field results indicate that eDNA is not equally effective for all species and was particularly ineffective in capturing the *Caulerpa* signal in the field using standard protocols. We demonstrate the value in benchmarking eDNA assays both in the lab and in the field prior to its deployment as a monitoring tool. Best practices in method validation should be adopted for all eDNA assays to ensure that results in the field, such as the negative results obtained in this study, are properly scrutinized and validated.

With any new methodology it is important to understand its strengths and limitations. This is especially important so that researcher and managers can maintain reasonable expectations when deploying novel approaches. There may be times when eDNA-based approaches cannot adequately replace traditional methods. Researchers must use caution and conduct rigorous validation of eDNA assays in the field and lab to understand the efficacy within a given system. Future limitations for the detection of *Caulerpa* eDNA may be ameliorated with the development of a more sensitive assay (perhaps targeting a chloroplast or mitochondrial marker gene) or the filtration of larger water volumes. Ultimately, we demonstrate the limitations of eDNA as a survey tool as it relates to the invasive algae *Caulerpa prolifera* and demonstrate the importance of contextualization and validation of eDNA assays for biomonitoring applications.

## Author Contribution Statement

Conceptualization: TW, ZG, ST, RE. Formal Analysis: TW, KL, ZG. Funding Acquisition: TW, ST, RE. Investigation: TW, KL. Methodology: TW, KL, ZG. Supervision: ST, RE. Writing - Original Draft Preparation: TW. Writing - Review and Editing: All authors.

## Data Archiving

Data from this study will be included in the supplemental file for this manuscript.

## Acknowledgements

Tanner Waters is supported by the UCLA Cota-Robles Fellowship, the National Science Foundation Graduate Research Fellowship, the Switzer Foundation Fellowship and the UCLA Center for Diverse Leadership in Science Fellowship. Environmental DNA work in the Eagle lab is supported by a grant from the David and Lucile Packard Foundation to the Center for Diverse Leadership in Science at UCLA. Robert Eagle also acknowledges support from the Pritzker Endowment to UCLA IoES. The authors acknowledge Dr. Jeana Drake and the Jacobs lab for the assistance in the use and access to the tanks used in the experiment. We thank Dr. Joshua Steele for his help in answering primer/probe design questions and Dr. Kim Parsons and Ana Ramón-Laca for their help in PCR troubleshooting. We also want to acknowledge Brian Owens, Christopher Potter, Terri Reeder, Keith Merkel, Dr. Robert Mooney, and others at California Department of Fish and Wildlife, the California Water Board, and Merkel & Associates for their help in identifying, surveying, and removing the Caulerpa from Newport Bay.

